# ICM cells can be partitioned robustly through transient synchronization using secreted FGF4

**DOI:** 10.1101/2020.12.15.422839

**Authors:** Xiaochan Xu, Ala Trusina, Kim Sneppen

**Affiliations:** Niels Bohr Institute, the Center for Models of Life, Copenhagen, 172100, Denmark

**Keywords:** Multipotent, genetic switch, bistability, bifurcation, FGF4 feedback, robust partitioning, regulatory logic, cell proportion, cell-cell communication

## Abstract

The differentiation of ICM cells into epiblast (EPI) and primitive endoderm (PE) is central in embryonic development. It is known that FGF4 signaling is important in this process, but it remains unclear how cells can be correctly partitioned. Here we model the NANOG-GATA6-FGF4 network, and test all 64 logical regulatory combinations for their ability to partition a group of cells. We found that nearly all the logic combinations allowed for correct partitioning, including a minimal network where self-activation of NANOG and GATA6 was inactivated. However such self-activation increased the robustness of the system. Furthermore, the model also captured the reported changes in cell proportions in response to FGF perturbations. This constrains the possible regulatory logic and predicts the presence of an “OR” gate in cell-cell communication. We repeatedly found that FGF4 coordinated the decision in two phases: A convergence and a bifurcation phase. First FGF4 negative feedback drives the cells to a balanced “battle” state where most cells have intermediate levels of both regulators, thus being double positive. Subsequent bifurcation happens at constant FGF4 level. Together our results suggest that the frequently observed state of multipotency during differentiation may be an emergent phenomenon resulting from inter-cellular negative feedbacks.

## Introduction

The early mammalian embryo consists of three distinct layers: trophectoderm (TE), epiblast (EPI) and primitive endoderm (PE). Both EPI lineage and PE lineage originate from inner cell mass (ICM). EPI cells give rise to the whole fetus while PE cells are limited to the yolk sac to support the growth of the fetus^1–3^. One recurrent feature during embryogenesis is that cells go through an intermediate state of multipotent progenitors. During ICM differentiation, these cells are referred to as double positive (DP) cells, with comparable levels of lineage specifiers, NANOG and GATA6. The expression levels of the specifiers start at low and initially increase in time, but later bifurcate into mutually exclusive states^4,5^, one with high NANOG and low GATA6 (EPI) and the other with low NANOG and high GATA6 (PE). It is elusive whether the DP cells are in a stable or transient state during the cell fates decision.

The differentiation is driven by a genetic switch in each cell, modulated by a cell-cell signaling factor (here FGF4, Figure 1A). FGF4 is secreted by NANOG-high cells and favors the opposite GATA6-high cell type^6,7^. Thereby a collection of cells can self-organize to the desired proportion of each cell type (EPI:PE ≈ 40%: 60%)^8^. The combination of local decision-making and external cell-cell communication is also found in other differentiation processes including the Notch signaling pathway^9^. However, it is currently unclear which components of the regulatory network (Figure 1A) are important for cells to self-organize into the right proportions.

**Figure 1.**
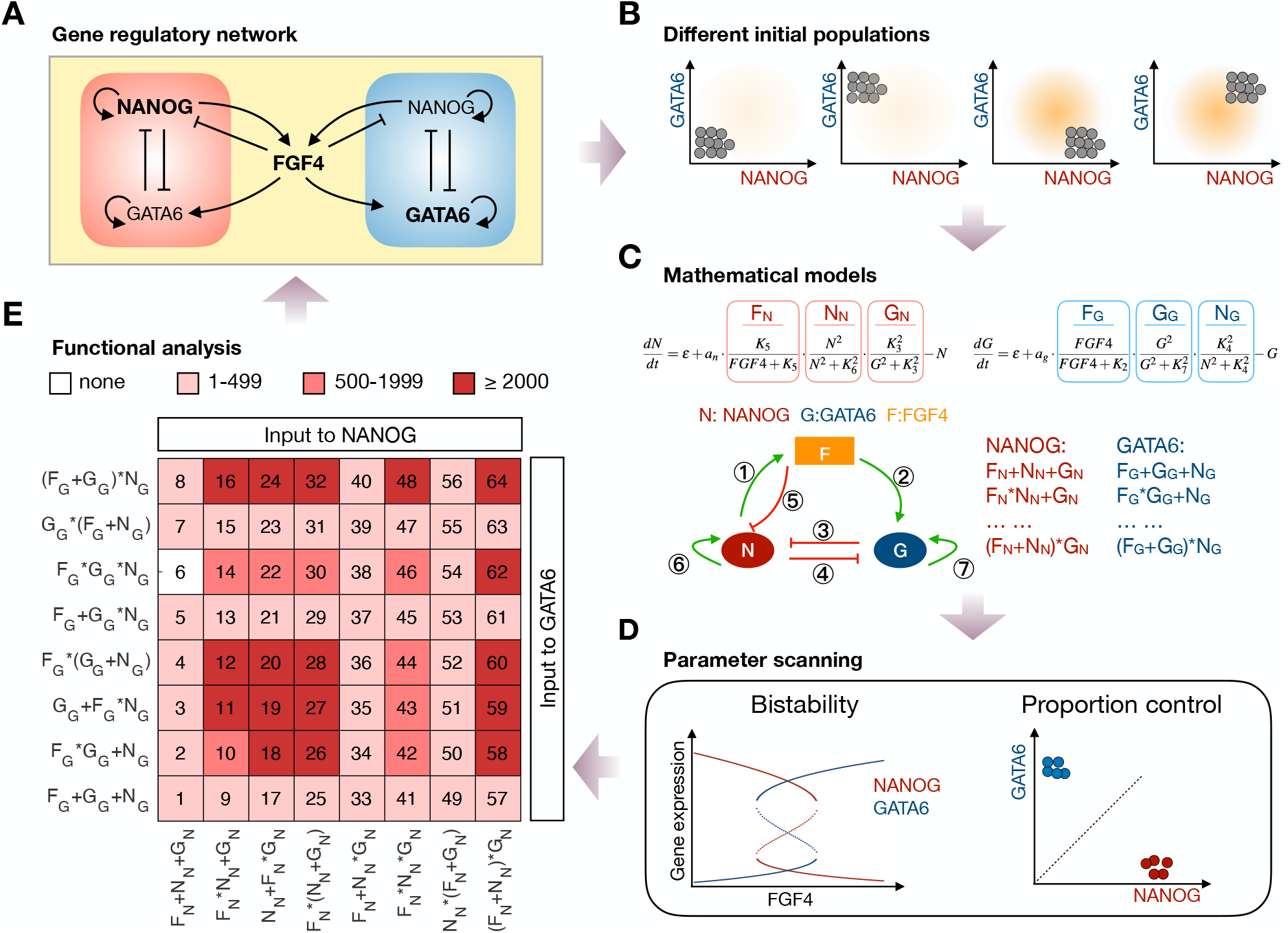
Theoretical framework for investigation of ICM differentiation. (A) Simplified gene regulatory network of NANOG, GATA6, and FGF4. (B)-(E) Schematic of the systematic analysis of the regulatory motifs for their ability to keep cell proportions. (B) Four different initial populations are used for parameter scanning (NANOG:GATA6 was low:low, low:high, high:low, or high:high). The background color indicates FGF4 levels, light yellow (low) and yellow (high). (C) For the original circuit architecture in (a), we test different logical combinations of “AND” and “OR” rules. *F_N_*, *N_N_* and *G_N_* represent the inputs to NANOG, while *F_G_*, *N_G_* and *G_G_* are inputs to GATA6. The *i_th_* edge (left) has the corresponding Hill constant *K_i_*. (D) Two criteria for identifying working parameters. First (left), two cell fates coexist with proper FGF4 level. Second (right): Starting with the initial conditions in (b), cells partition into the right proportions at steady state. (E) The ability of the different logical combinations to produce correct cell proportion. 14,641 parameters were scanned for each logical model. The shades of red color show the numbers of working parameters for each model. The numbers in the matrix refer to model identities, each with unique logical combinations (specified to the left and at the bottom).

Recently, using quantitative experiments and a simple mathematical model, Saiz et al.^10^ has proposed that FGF4-mediated cell-cell communication is sufficient for the robust maintenance of cell proportions. At the same time, there is an alternative scenario where cells, relying on cell-intrinsic genetic switch alone, can bifurcate into the final fates due to stochastic fluctuations in protein concentrations. Furthermore, it is unclear if the mixed, DP, state is an emergent property of a robust cell partitioning, or a simple coincidence.

Here we systematically study all 64 combinations of input logics for the regulatory network that determine the partition of ICM cells. While most logical combinations may work, we will see that it is much easier to find working parameters for some architectures than others. In particular, our investigations will point to the added robustness if FGF4 both inhibits NANOG and activates GATA6, and we will see that self-activation of NANOG and GATA6 also help. Universally we will see that the DP state emerges from the collective cell behavior mediated by the lateral inhibition from FGF4. Overall our simulations robustly generate a differentiation dynamic where an initial convergence among the ICM cells is followed by a subsequent divergence into the two final cell types.

## Results

### Mathematical model of the early embryo development

ICM differentiates at E3.0~E3.5 when the embryo has about 32 ICM cells. We simulated a system with 36 cells. It has been conclusively shown that the differentiation depends on the core gene regulatory network of the lineage specifiers, NANOG and GATA6, and the intercellular FGF4 signaling^6,7^. NANOG and GATA6 mutually inhibit each other within the cell. The FGF4 is mainly secreted by NANOG-high cells and subsequently influences gene dynamics in the neighboring cells (Figure1A). Both FGFR1 and FGFR2 are the crucial receptors for FGF4 signaling. However, FGFR2 is restricted in PE cells^11^ while all the ICM cells respond to FGF4 through FGFR1^12,13^. This suggests that the FGF4 signaling is mainly through FGFR1 in the early cell fate separation.

The FGF4-mediated gene regulation involves several steps between the external receptor and the final gene expression^14,15^, and the gene regulatory network in Figure 1A simplified this pathway into one step. We used Hill functions to describe the simplified gene regulation. The inhibition from regulator *X* was characterized as *K^h^*/(*X^h^* + *K^h^*) and activation as *X^h^/*(*X^h^* + *K^h^*), *K* is the dissociation constant and *h* is the Hill coefficient. The Hill coefficients of equation (1–2) were fixed to 2. The one of equation (3) was set to 5 (*h* = 5) in the discussion of different logics since we found the model is more robust with high Hill coefficient as input to FGF4. *h* is only referred to the Hill coefficient in equation (3) hereafter. There were 9 parameters for the regulatory network (Figure 1C): *K*_1_, *K*_2_,…*K*_7_ were dissociation constants for each regulatory, and *a_n_, a_g_* were factors for scaling the expression levels of NANOG and GATA6 to around 1. Four free parameters (*K*_2_, *K*_5_, *K*_6_ and *K*_7_) varying log equidistantly from 0.05 to 5 (see methods).

For example, if all the inputs to both NANOG and GATA6 followed the “AND” logic, the model is:

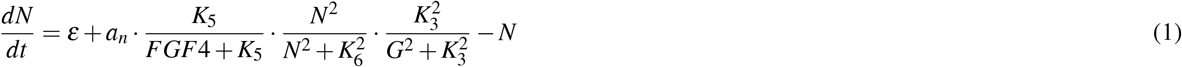

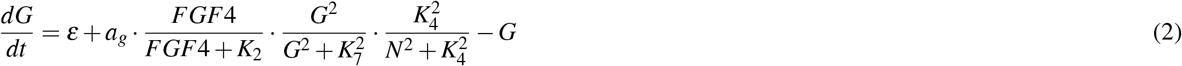

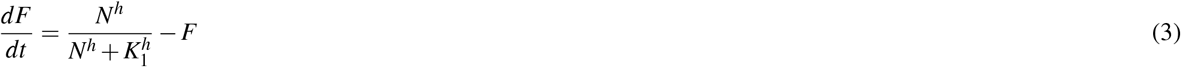

Here *N*: NANOG, *G*: GATA6, *F*: FGF4 produced by the cell, *FGF*4: the FGF4 signal the cell had received and was calculated as the average of FGF4 produced by itself and its neighbours. This model was labeled as *F_N_* * *N_N_* * *G_N_*; *F_G_* * *G_G_* * *N_G_* (logic 46 in Figure 1E).

Heterogeneity between individual cells, especially the variable NANOG expression, has been observed in transcription and dynamics in the early development at single cell resolution^16,17^. Despite this, embryos can avoid fluctuations and robustly maintain a normal phenotype with correct cell proportions. The heterogeneity between the cells was simulated by adding noise to gene expression in the initial population. A recent study shows the embryo can restore lineage composition even if one of the lineages has been ablated^10^. Here, we chose 4 extremely different initial cell populations (Figure 1B) to validate whether the model could produce the right cell proportion under different situations. These four initial populations simulate double negative cells at the beginning of the development, all GATA6-high cells, all NANOG-high cells, and all double positive cells.

The knowledge about the structure of the gene regulatory network is not sufficient to map the network to the equations. When a gene is regulated by several factors one has to decide on what logic to use, one needs to specify whether the regulatory inputs should be added or multiplied, reflecting a logical “OR” or logical “AND” functions (Figure 1C). For the core regulatory network in Figure 1A, there were 64 different logical combinations (summarized in Figure 1E).

Given the small number of cells in the embryo, we first assumed that FGF rapidly diffused across all cells, and the spatial structure could be omitted. The average level of FGF4 in the system was used as an input for the cells’ response to external FGF4 signaling. This generalized model (referred to as “Global model” hereafter) allowed us to theoretically analyze whether our method was applicable and which logical combinations were functional.

FGF4 is required for the co-existence of NANOG-high cell and GATA6-high cell in the embryo^6,18,19^. FGF4 null mutant embryo entirely lacks GATA6-high cell while extra external FGF4 in the medium can rescue the mutant phenotype. For the model to be considered functional, simulation results had to be consistent with the following experimental observations (Figure 1D, left).

- A given cell could result in either a stable state with high NANOG and low GATA6, or in a state with low NANOG and high GATA6.
- No GATA6-high cells when FGF4 was fixed to low, and no NANOG-high cells if FGF4 was fixed to high. The cell could become a NANOG-high or GATA6-high cell if FGF4 was fixed to an intermediate level.

Furthermore, we tested all the models on its ability to partition cells in two distinct cell-fate populations of equal size. More specifically, we required that a functional model should also differentiate all of the 36 cells to either NANOG-high cell or GATA6-high cell, and the percentage of each cell fate was limited to 50% ± 15% regardless of the initial conditions (Figure 1D, right).

Our method recapitulated the features of the core gene regulatory network, since all the 64 logical models could give bistability and 63 of them could partition ICM correctly into the two cell types with working parameters (Figure 1E). But the logical models had different robustness of the parameters. Models with logics including the inputs to NANOG as *F_N_* + *N_N_* + *G_N_*, *F_N_* + *N_N_* * *G_N_*, *N_N_* * (*F_N_* + *G_N_*) or similar logics for the inputs to GATA6 had less working parameters than the others. 17 logical models had more than 2,000 (total 14,641) parameters making them functional for keeping cell proportions (Figure 1E, Figure S1).

### The simplest topologies capable of producing correct cell proportion

Having confirmed that the core gene regulatory network, compiled from experimental observations, could maintain cell proportions, we asked if it could be further simplified while maintaining the functionality. For this we first quantified which of the regulations and the respective parameters were important for the bistability and keeping cell proportions. We looked into the working parameters and found different logical models had different preferred parameter distributions (Fig 2A) except that the core of the genetic switch had nearly symmetric NANOG:GATA6 repression (same strengths of *K*_3_ and *K*_4_ when measured in units of the corresponding maximal expression). Some of the working parameters were small (= 0.05) or large (= 5), indicating that the network still could work for keeping cell proportion even though the corresponding regulatory link was abolished. That is, for *K* ≪ *G* ~ 1 then activation *G/*(*G* + *K*) ~ 1 whereas repression *K/*(*F* + *K*) ~ 1 when *K ≫ F* ~ 1.

**Figure 2.**
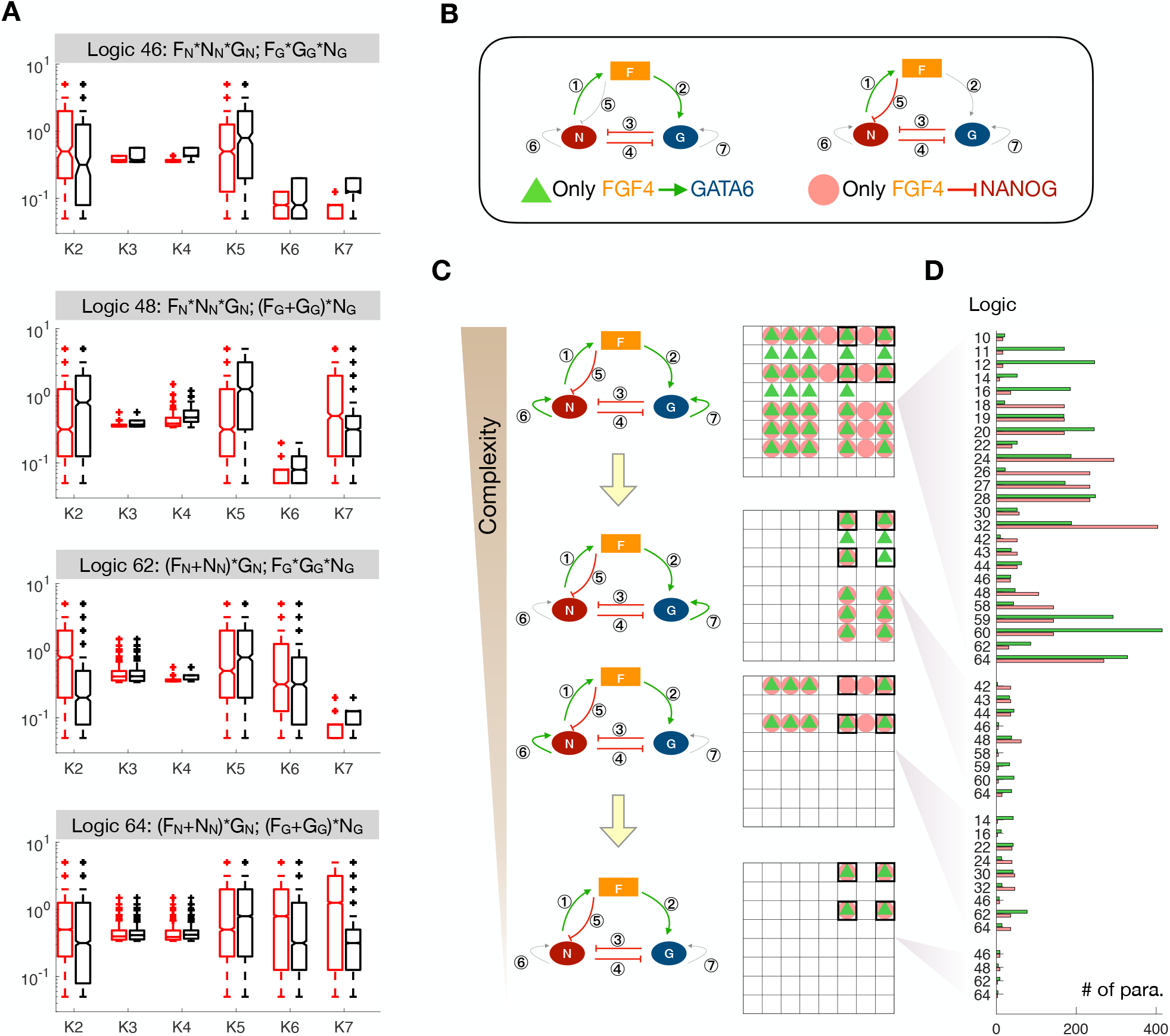
The simplest topologies partition correctly. (A) Statistics of the working parameters for the models with logic 46, 48, 62 or 64. Black: only obtained bistability, red: obtained bistability and partitioned correctly into 50 ± 15%. +, values > 1.5 times times the interquartile range. (B) The simplest topologies that can partition correctly. Left, FGF only activates GATA6; Right, FGF4 only inhibits NANOG. (C) Self-activation of NANOG or GATA6 increases the number of working parameters and logics. Grey links mark abolished edges. The matrix to the right corresponds to Figure 1e. Green triangles and pink circles mark FGF regulations specified in (B). Black rectangles mark the logics shown in (A). (D) The numbers of working parameters with only activation of GATA6 or with only inhibition of NANOG are compared. We also show logics which generate working parameters when one of the links from FGF4 is abolished. Green and pink colors correspond to the FGF regulation shown in the left and right panel of (B).

For example, Models with logic 46, 48 or 62 have weak self-activation links of NANOG or GATA6 with low values of *K*_6_ and *K*_7_ (≪ 0.5). It suggested that these self-activation links were not necessary to direct differentiation into proper cell proportions.

By abolishing the regulatory edges of the network one by one, we found that proper cell proportion could be obtained by the simplest topology with FGF4 only activating GATA6 or with FGF4 only inhibiting NANOG (Figure 2B, C). The fact that there are many more working parameters with both the activities of FGF4 signaling in Figure 2D suggested that the action of FGF4 on both NANOG and GATA6 contributes to correct cell partitioning.

The importance of the FGF4 dual role for the correct partitioning was also reflected in the parameter scans that persistently selected high *K*_2_ and low *K*_5_ (Figure 2A). It reflected that correct cell partitioning required both regulations to be sensitive to FGF4 signaling (activation ∝ *F/*(*F* + *K*_2_) ∝ *F*, inhibition ∝ *K*_5_*/*(*F* + *K*_5_) ∝ */F*).

The complete network performed best at controlling proportions (Figure 2C,D). Abolishing NANOG or GATA6 self-activation reduced network complexity (both the number of possible logics and the number of free parameters). However, even though these two edges were not necessary, they increased the robustness of the correct cell partitioning in the sense that they increased the chance that a random parameter worked by an order of magnitude. And they did that in spite of the fact that more complex networks involve more parameters and thus should be expected to have large space for failure.

### Convergence and bifurcation phases of the FGF4-mediated cell-fate decision

The requirements for the functional models included both complete cell-fate separation (each cell achieved either NANOG-high cell fate or GATA6-high cell fate) and maintenance of the right proportions in the range 50% ± 15% starting from 4 different initial conditions (Figure 1B).

By requiring models to partition correctly starting from either of the four initial conditions, we imposed more stringent constraints than those observed in living embryos (In an embryo, cells start with low levels of NANOG and GATA6). Thus the criterion for acceptance of a network motif included a requirement of being robust against variations of initial conditions.

Figure 3A, B illustrates the dynamics of NANOG and GATA6 for a collection of cells using working parameters for the models with logic 46 (pure “AND”) and logic 16 (mixed) from Figure 1. Strikingly, the individual cells first tended to converge in gene expression, then later to partition into the two cell types. It was seen in Figure 3 where the expressions of NANOG (red curves) of all individual cells converged to a certain level and at the same time the expressions of GATA6 also converged. Thus the cells dynamically self-organized towards the saddle point of an individual cell at an externally fixed FGF4. Figure 3C, D, show that the difference in expression level for individual cells to this saddle point is smallest at this time point. After the convergence, the FGF4’s level (yellow curves) remained nearly constant while gene expression in individual cells differentiated into either a NANOG dominated state or a GATA6 dominated state (Figure 3A, B).

**Figure 3.**
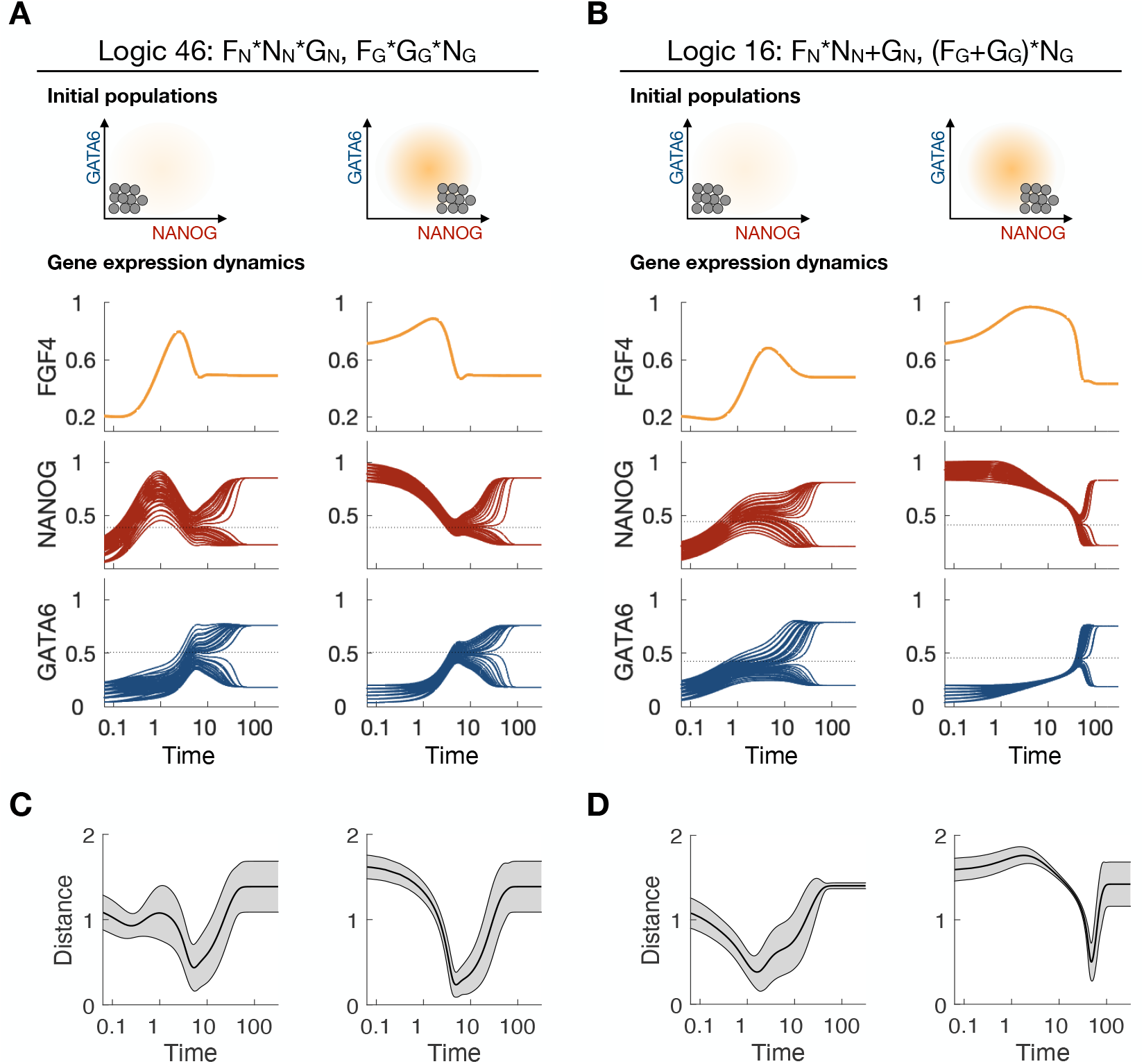
Cells governed by functional models converge to the “battle” before committing to different cell fates. (A)-(B) The dynamics of the mean FGF4 (yellow, the second row) of the 36 cells, NANOG (red) and GATA6 (blue) in each cell with different initial conditions (first row) for models with logic 46 (A) or logic 16 (B). The dashed lines show the levels at the “battle”. (C)-(D) Evolution of the difference between the instant gene expression and the levels at the “battle” point for the 36 cells. Black marks the mean while grey shows the standard deviation. The values of NANOG, GATA6 and FGF4 is each normalized by their respective level at the “battle state” when calculating the distance. The shown distance is calculated as the sum of the normalized mean square differences for the NANOG, GATA6 and FGF4 contribution of each cell.

We refer to this convergence-divergence state of the cells as the “battle”. The whole differentiation process was divided into two phases: the convergence phase and bifurcation phase.

To test if our results are sensitive to the assumption of the global FGF signaling (“Global model”), we have run the simulations where FGF signaling was short-range, only to the neighboring cells. In the following we refer to it as “Local model”. The difference between the “Global model” and “Local model” was that the cells only received FGF4 signaling from its neighbors. To model neighbor interaction, the cells’ positions in 3D and its nearest neighbors were assigned as described in methods section.

We found that this did not change the results. The logics that worked well without locality also worked well with local interactions, and a relatively large fraction of the working parameters scanned from the “Global model” also worked with the “Local model” (Figure S1). In both cases the “salt-and-pepper” pattern was seen (Figure 4A, B), and the gene expression dynamics exhibited the convergence phase with subsequent bifurcation (Figure4C).

**Figure 4.**
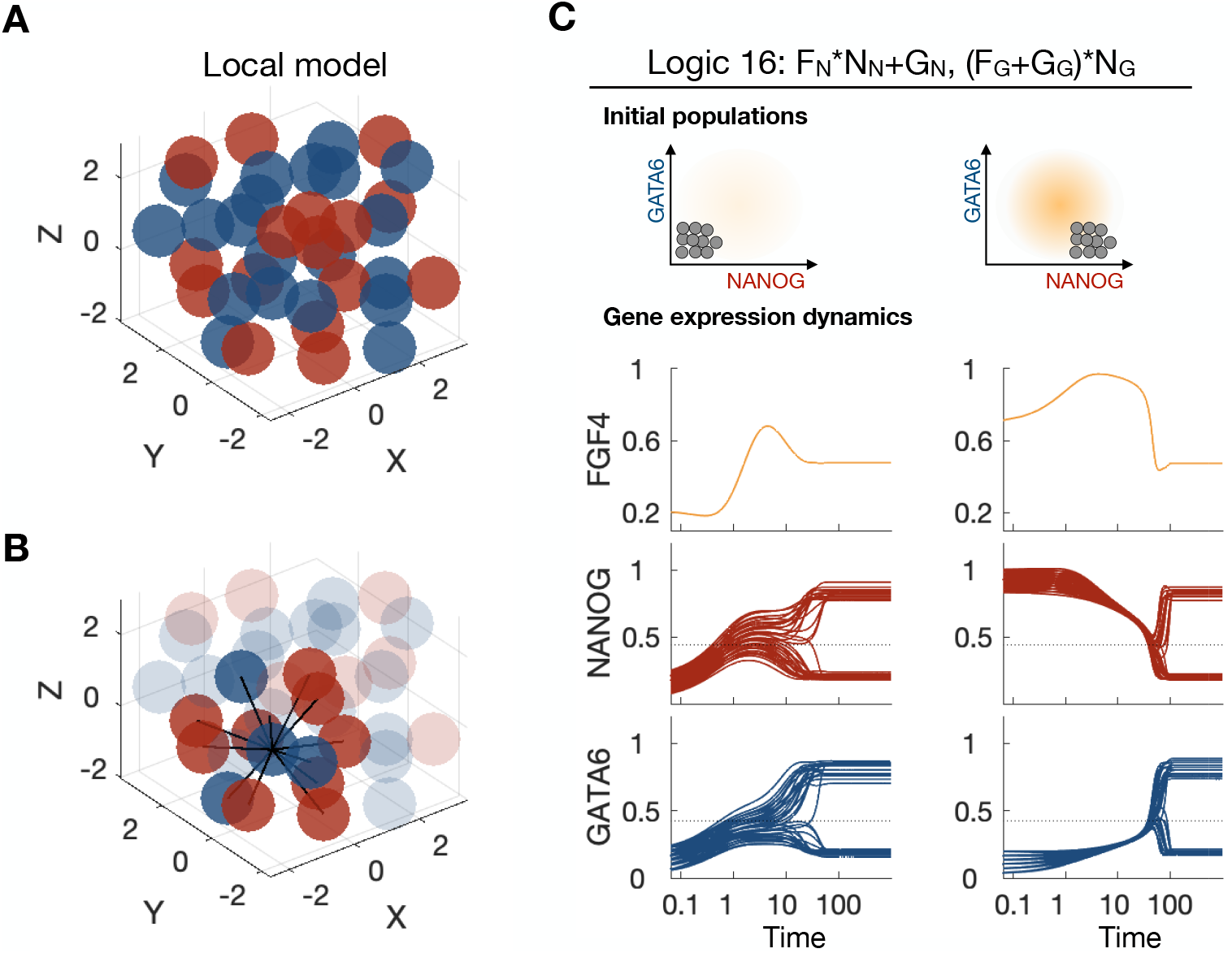
A “Local model” of the ICM differentiation. (A) Simulated “salt-and-pepper” pattern of differentiated ICM cells with the “Local model”. Blue: GATA6-high cells, red: NANOG-high cells. The positions of the cells were determined as described in methods and fixed during the time evolution of the differentiation process. (B) Example of one cell and its neighbors connected by black lines in the simulation with the “Local model”. (C) The gene expression dynamics of the mean FGF4 (yellow) of the 36 cells, NANOG (red) and GATA6 (blue) in each cell with different initial conditions. (A)-(C) Show results from the “Local model” using logic 16.

### “OR” logic can capture the changes in cell proportions with increasing FGF4

In experiments where embryos are cultured in a medium with exogenous FGF4, both the wild type embryos (Wt) and FGF4 homozygous embryos (FGF4−/−) have an increased proportion of SOX17+ cells^7^ (Figure 5A). Our “Local model” with “OR” logic can recapitulate this result, as the proportion of GATA6-high cells increased with increasing FGF4 dose, in line with experimental data (Figure 5C,D,F,G).

**Figure 5.**
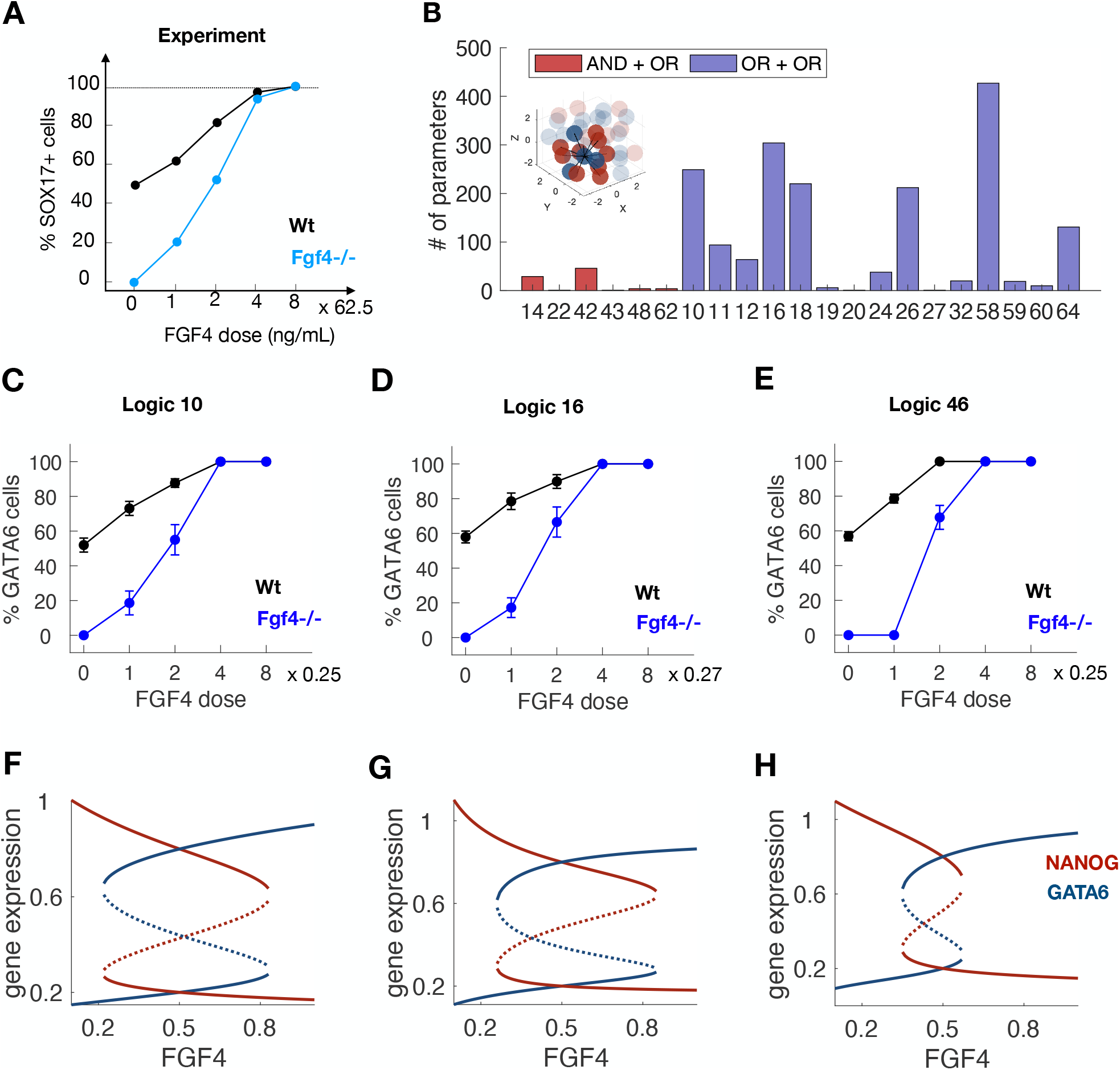
“OR” logic is required for the FGF dose-response. (A) The response of the cultured embryos to exogenous FGF4^7^. Black: Wild type embryos. Blue: Fgf4−/− embryos. The dots marked the mean proportion of GATA6-high cells. (B) The numbers of parameters keeping both cell proportion and soft dependence with different logics when applying the “Local model”. (C)-(E) Simulations of FGF4 dose-response with different logics. Black: Wild type embryos. Blue: Fgf4−/− embryos. Error bars show the simulated embryo to embryo variation in 20 repeats. (F)-(H) Bifurcation diagrams of the gene expression with different exogenous FGF4 levels using corresponding parameters in (C)-(E). Blue: GATA6, red: NANOG. Solid curves: stable steady state, dash curves: unstable steady state.

The working parameters for logical 46 (pure “AND”) exhibited a very high sensitivity to FGF4. Thus a small change in FGF4 induced the change from all NANOG-high cells to all GATA6-high cells (Figure 5E, H). This disagreed with the experiments where the shift from one to the other cell type happens over at least a four-fold change in external FGF4^7^. When one of the “AND” logic was substituted to an “OR”, it became much easier to find parameters that give a larger FGF4 response range. Such larger range is equivalent to a softer dependence quantified by the maximum change in cell proportion when external FGF4 was changed by factor 2. The response range was further improved with two “OR” logics (Figure 5B). Thus we predicted that the soft dependence of FGF4 from the FGF-titration experiments is governed by an “OR” logic between the regulatory inputs to NANOG. The examples of the simulation of FGF4 mutant with logic 10, 16 (Figure 5C, D, F, G) and logic 14, 42 and 64 (Figure S2) were shown.

The Hill coefficient *h* in equation (3) had effects on the compromise between the soft dependence and cell proportion control. When *h* decreased from 5 to 3, there were fewer working parameters for the working logics (Figure S2). And if *h* = 1, no parameters worked with any model. The larger *h* guaranteed the cells needed a certain level of NANOG to produce FGF4. The GATA6-high cells only existed when NANOG-high cells existed in the system to provide enough FGF4, which made the FGF4 level in the multi-cell system sensitive to cell proportions.

### The negative feedback on NANOG through FGF4 drives the cells to the “battle” state

Each cell can have 3 steady states, two stable steady states (NANOG-high or GATA6-high) and one unstable steady state (intermediate NANOG and GATA6), with intermediate FGF4 in the system. The stability of the steady states of the system depends on each cell’s stability. The steady state is unstable even only one cell is at the unstable steady state. The cells will be identical if all of them are at the unstable steady state, and the system is at a special unstable steady state (Figure S3A). The system is at a stable steady state if all of the cells are at stable steady states with current FGF4 in the system. The different proportions of NANOG-high cells and GATA6-high cells demand the different FGF4 levels in the system and will produce different amounts of FGF4 (Figure S3B). The system can only maintain stable when the FGF4 produced by the cells balances the FGF4 demanded by the cell proportions. The system has other unstable steady states. The FGF4 produced by the cell at the unstable-steady state is higher than at the stable GATA6-high steady state and lower than at the stable NANOG-high steady state. Thus these unstable steady states are between two arbitrary stable steady states.

Most intriguingly, even though the FGF4 level in the system changes dynamically, the shape of the stable manifold regarding the single-cell system does not change much. All of the trajectories in the gene expression space converge to the stable manifold first and move along the manifold (6A-D, Figure S3 C-E). When the varied FGF4 changes the number and state of the attractors on the manifold, the cells close to or on the manifold will change their directions synchronously if only one attractor exists and will separate into two groups if two attractors exist. Before the bifurcation, the cells are most likely to behave similarly and move towards the special unstable steady state in Figure S3A.

**Figure 6.**
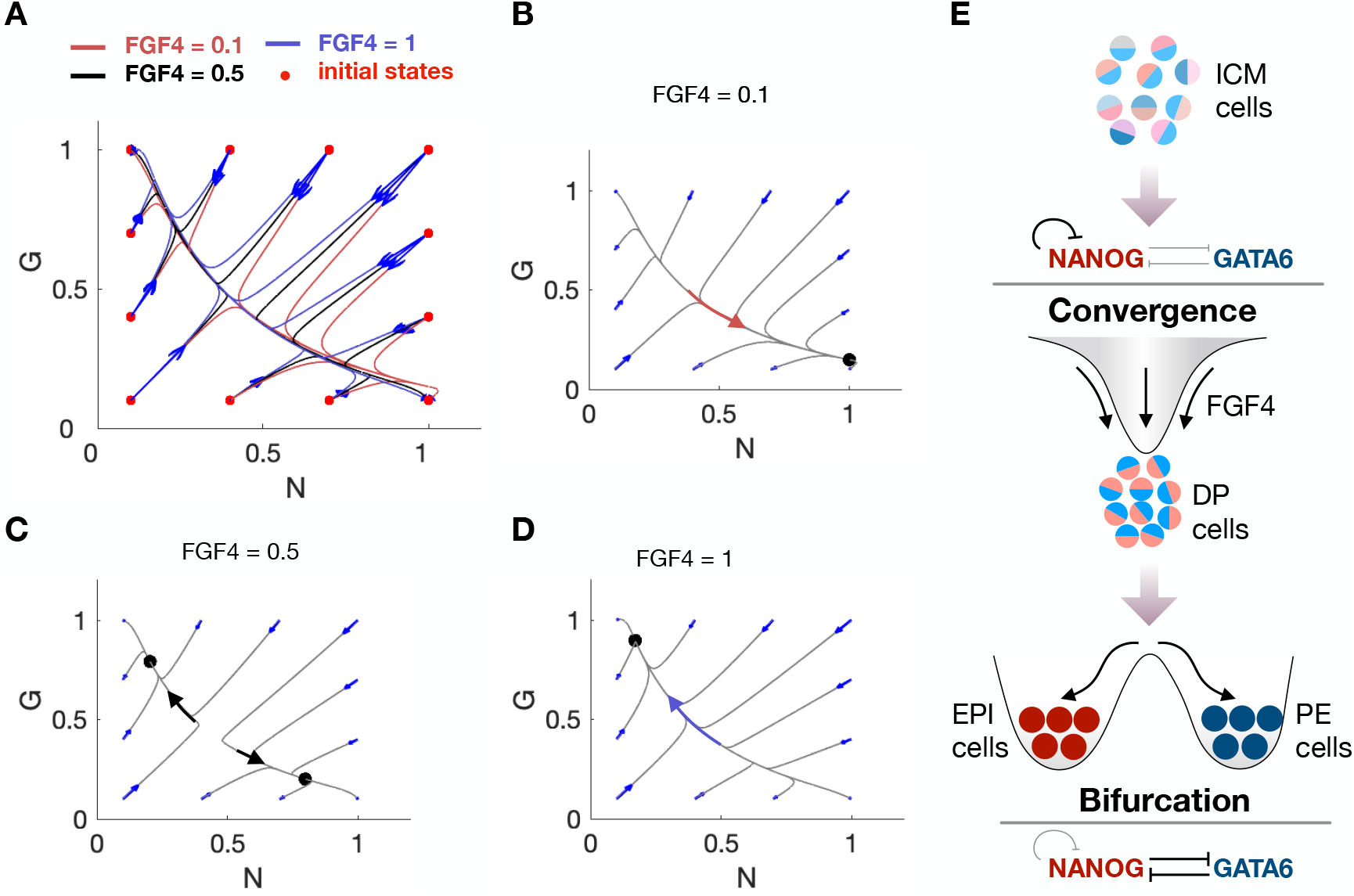
Two phases model of how FGF4 helps partition ICM cells correctly. (A) Trajectories of cells converges to the unstable manifold of the system. Red dots, the different initial states of one cell; curves, trajectories of the gene dynamics starting at the corresponding initial state with different FGF4 level in the system; blue arrows, directions of the trajectories. (B) Trajectories starting at different initial states when the FGF4 in the system is low. The cell converges to the manifold and then moves along the manifold towards the stable state where NANOG is high and GATA6 is low. (C) Trajectories starting at different initial states when the FGF4 in the system is intermediate. The cell converges to the manifold and then moves along the manifold towards the stable states where either NANOG is high or GATA6 is high. (D) Trajectories starting at different initial states when the FGF4 in the system is high. The cell converges to the manifold and then moves along the manifold towards the stable state where NANOG is low and GATA6 is high. (E) Two phases model of how FGF4 helps partition ICM cells correctly. The ICM cells converge towards the unstable steady state (showing as DP cells) of the system due to the negative feedback of FGF4, then bifurcate to different cell fates, EPI cells or PE cells, due to the mutual inhibition of NANOG and GATA6. The black dots in (B)-(D): the attractors of the expression of NANOG and GATA6 regarding the different levels of FGF4 in the system; Colored arrows on the manifold, directions when cell moves along the manifold.

This systematic property suppressed the variation between cells and lead to convergence to the DP positive state. The system can achieve the unstable steady state only when the cells have identical initial conditions, which is impossible in reality. Thus bifurcation of gene expression dynamics happens after cells get close to the unstable steady state.

Overall the robust finding of our modeling is the convergence to a “battle state" followed by a divergence of individual cells into the two cell types (Figure 6E). The early development is driven by FGF4 mediated negative feedback on expression of NANOG from all cells. This feedback leads to an unstable intermediate expression of NANOG and an associated intermediate level of GATA6, thus identifying as the DP state. The cells are collectively driven towards this “battle” state, but when reaching this state they each become exposed to the double positive feedback between NANOG and GATA6. In the most primitive version of the model, this leads to a random partitioning of cells into the two cell fates. Thanks to negative feedback through FGF4 eventual mistakes in the partitioning can be partly corrected such as an excess of NANOG cells would increase the likelihood for undecided cells to move towards the GATA6-high state.

Noticeably, the cell partitioning is fastest when the negative feedback through FGF4 is not too slow. The system will move to the “battle" directly since FGF4 will change immediately with NANOG (Figure S4). Conversely, if FGF4 is slowly degraded, the system could initiate damped oscillations around the “battle state", which in turn would delay the cell partitioning (Figure S5). Thus our study identified the DP state as a “battle” state. It suggests that a self-organization process mediated by cell-cell communication obtained the DP state and allows subsequent cell fate partitioning through positive feedback inside each cell.

## Discussion

During development cell populations should organize to the right proportions of cell types and dynamically correct to situations where the proportions are perturbed. Organs like heart^20–22^, liver^23,24^, and pancreas^25,26^ can regenerate and restore their function when impaired. Cell-cell communication has been suggested to be an important part of such repair^27,28^.

Our results suggest that a differentiation trajectory, where cells go through a “battle” state before bifurcating, may be a generic consequence of negative feedback through cell-cell communication. This behaviour is typical when cells start from states with similar gene expressions. This overall dynamics agrees well with the development of embryonic stem cells, where progenitor cells start with a low level of transcription factors associated with the future fates.

An extensive sampling of possible parameters for the ICM differentiation network showed that precise and robust cell partitioning could be obtained by many combinations of “AND” and “OR” gates to each of the regulatory genes. However, the experimental observation of the relatively soft sensitivity to external FGF4 signals in ICM development constrains the logical input to including “OR” logic between the FGF4 signal and self-activation of the central genes. Cell partitioning was not performed better or more precisely by the partial “OR” logic, and our model does not explain why the system appears to use “soft” decision logic. Perhaps it is due to the cooperation between the FGF4 signaling and other signaling pathways involved in the differentiation outside the system considered here. Importantly our findings are valid for both a local cell-cell interaction, where cell signals to its nearest neighbors and global interactions, where singling molecules diffuse over several cells. If signaling is truly local, then cells at the periphery of the embryo will have fewer NANOG-high cell neighbors, and thus tend to become NANOG-high cell. When dependence of the exact level of FGF4 is softened this effect is weaker, and thereby it is easier to obtain the correct cell spatial partitioning of NANOG and GATA6-high cells after the “salt-and-pepper” state.

In this paper, our models mainly focus on cell-cell communication, while the real biological system is much more complex than considered. Besides cell-cell communication, other phenomena like cell death^29^ and polarity^30^ also play important roles in ICM differentiation. Our models can be extended to study those aspects. Some cells may differentiate later than the others or even be undifferentiated until late. It is probably due to the locality of FGF4 signaling, which could be seen from our “Local model”. Or the system has not achieved a stable steady state. Some cells happen to stop at the intermediate unstable steady state.

We predicted that the models with the “OR” logic of the gene regulation inputs to NANOG or GATA6 are closer to reality. To narrow down the specific logics which the ICM adapts is possible by ruling out the models with other experiments besides the mentioned FGF4 soft dependence.

In our work, we persistently found that FGF4 controlled ICM cell partitioning in two dynamic phases. In the early “convergence” phase, the system organized the cells into a collective state with FGF4’s feedback. In the subsequent differentiation phase, the FGF4 was maintained at the same level, while the cells diverge away from the “battle”. In the language of epigenetic landscapes of Waddington, our FGF4 first deforms the landscape into a funnel, and when FGF4 has reached an intermediate level, the funnel is divided by a mountain ridge formed by the positive feedback inside each cell. We believe our predictions will inspire further research into the cell proportion maintenance during both the ICM differentiation and other developmental systems where cell-cell interactions may regulate cell proportions.

## Methods

### Identifying free parameters and parameter constraints

The list of the constraints and their rationalization is summarized in Table 1. All of the parameters used for the main figures are summarized in supplementary materials.

For simplicity we set the half-lives for GATA6 and NANOG to be the same and the time to be unitless, rescaled by the NANOG/GATA6 half-life. The full network in Figure 1 had 9 free parameters: *K*_1_, *K*_2_,…*K*_7_, *a_n_*, *a_g_* (*∊* was fixed to 0.05).

**Table 1.**
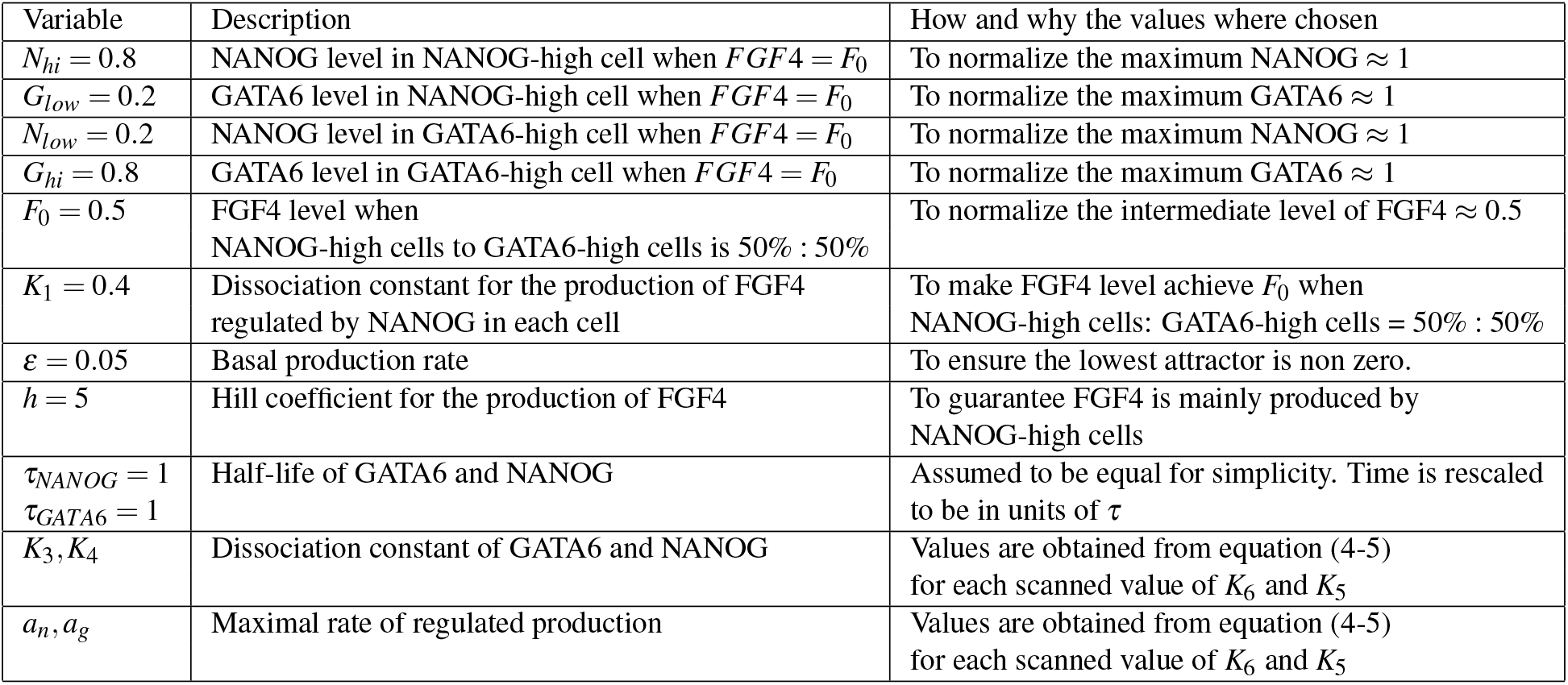
Imposed constraints. The table show that a cell that is differentiated into NANOG dominated state at *FGF*4 = 0.5 will have *NANOG* = 0.8, while a cell in GATA6 dominated state will have *NANOG* = 0.2 (and reversed for GATA6). These fixed levels of NANOG and GATA6 reduce the number of free parameters, and make it possible to compare the binding constants *K* in Figure 1 directly with the characteristic scale of 1.

We limited the ranges of these parameters to where they have regulatory effects, given that the maximal expression level of NANOG, GATA6, and FGF4 is around 1. By setting up proper gene expression levels and the fractions of the final cell fates, we could reduce the number of free parameters. We assumed that the stable state of the NANOG-high cell was (*N_hi_, G_low_*), the GATA6-high cell was (*N_low_, G_hi_*) and when each cell fate composed 50% of the population, the *FGF*4 = *F*_0_. *N_hi_* and *G_hi_* are levels of NANOG and GATA6 when these two genes get to the high stable steady state, *N_low_* and *G_low_* are levels of NANOG and GATA6 when these two genes get to the low stable steady state.

If

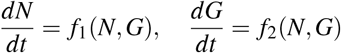

we had *f*_1_(*N_hi_*;*G_low_*) = 0; *f*_1_(*N_low_*;*G_hi_*) = 0; and *f*_2_(*N_hi_*;*G_low_*) = 0; *f*_1_(*N_low_*;*G_hi_*) = 0;

Taking the all “AND” logic model (logic 46) as an example, from equation (1) we had:

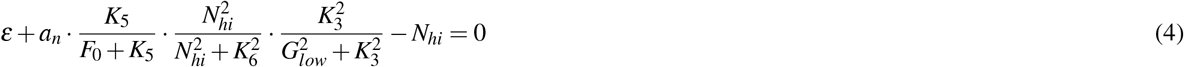

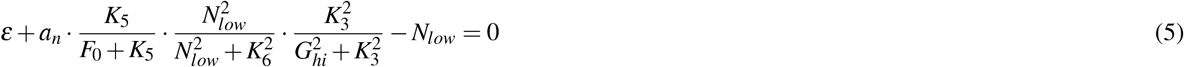

*K*_3_ and *a_n_* were solved through equation (4) and (5) when *K*_6_ and *K*_5_ varied log equidistantly and *N_hi_*, *N_low_*, *G_hi_*, *G_low_*, *F*_0_ and ɛ were fixed. Similarly, *K*_4_ and *a_g_* were solved when *K*_7_ and *K*_2_ varied log equidistantly.

Considering steady state level of FGF4 from equation (3), each NANOG-high cell contributes with

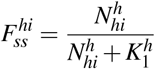

and each GATA6-high cell contributes with

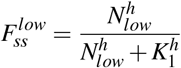

Assuming that NANOG-high cells: GATA6-high cells = 50%: 50% when *FGF*4 = *F*0, we got the FGF4 level at the steady state as

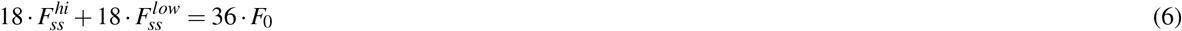

and solved *K*_1_ through equation (6). All the fixed values are summarized in Table 1.

With this strategy, only 4 free parameters (*K*_2_, *K*_5_, *K*_6_ and *K*_7_) needed to be scanned. For each set of *K*_2_, *K*_5_, *K*_6_ and *K*_7_, we could get *K*_3_, *a_n_* and from fixing *N_hi_* and *N_low_*, *K*_4_ and *a_g_* from fixing *G_hi_* and *G_low_*. We get *K*_1_ from the fixed *N_hi_* and *N_low_* when cells approach 50%: 50% partition. The same principle was applied to other models.

### Testing for bistability

All the fixed points of each model were solved with different parameters. For each fixed point, the Jacobi matrix was used to analyze the type and stability of it.
if

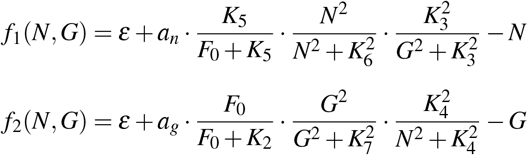

and at each fixed point (*N*\*, *G*\*)

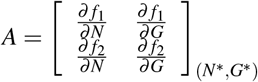

had Δ = *det*(*A*)*, τ* = *trace*(*A*), we demanded Δ < 0 for the saddle point, Δ > 0, *τ* < 0*, τ*^2^ - 4Δ > 0 for the stable node. The parameters in a specific logic had (*N_hi_, G_low_*) and (*N_low_, G_hi_*) as two stable nodes and one saddle point were chosen for the following analysis.

### Initial conditions

We used 4 initial distributions of NANOG, GATA6 and FGF4 to test which parameters that provide correct partitioning into the two cell types. For each initial condition, the values of NANOG and GATA6 for the 36 cells are uniformly distributed in an interval of length 0.2. Low expression values selected in the interval [0, 0.2] and high expression distribution was in the interval [0.8, 1]. Each test consisted of simulations from 4 sets of initial conditions, reflecting our high demand for robustness. To simulate the experimental data for the FGF4 dose-response in Figure 5a, both the initial values of NANOG and GATA6 were randomly chosen in the interval [0.05, 0.45].

In nearly all simulations we set the initial contribution to FGF4 for each cell *i* to *N_i_/*(*N_i_* + *K*_1_) (wild type simulations). The only exception was in simulating the FGF4−/− phenotype where initial FGF4 was set to zero.

### Parameter ranges of active vs. inactive regulations

The parameter restrictions (Table 1) allowed us to directly evaluate the impact of a specific regulatory link from its *K*-value. Thus an activating regulation with *K* ≪ 0.5 is always completely “on” (*X/*(*X* + *K*) ~ 1), whereas a repressing link with *K* ≫ 0.5 correspond to the complete absence of repression (*K/*(*X* + *K*) ~ 1). When exploring the minimal topology of the gene regulatory network, the parameters *K*_2_, *K*_6_, *K*_7_ were set to 0 and *K*_5_ was set to 100. Thereby the corresponding regulations were effectively abandoned.

### “Local model”

The “Local model” implemented the same regulations as the standard “Global model”, but constrained the interactions in space such that each cell only sensed the mean FGF4 contribution from its neighbors defined as in^31^. Each logic was tested with all the same parameters as in the “Global model”.

To find cell positions and neighbor cells, cells were simulated as interacting particles. Thus, starting from a random configuration, they moved to their final locations based on the repulsive and attractive forces. The potential between cell *i* and *j*, separated by distance *r_i j_* was defined as *V_i j_* = *e*^−*r*_i j_^ - e^−*r*_*i j*_/5^. The movement of cell *i* at each time point was decided by the sum of the potential between all its neighbors. A three-dimensional interaction network of the cells was constructed from the final locations.

The models were solved by a standard MATLAB (R2019a, 64-bit) ODE solver (ode45). The parameters used in the figures can be found in supplementary materials.

## Acknowledgements

This project has received funding from the European Research Council (ERC) under the European Union’s Horizon 2020 research and innovation program under grant agreement No 740704 and Danish National Research Foundation (grant number: DNRF116).

## Author contributions

Xiaochan Xu, Ala Trusina and Kim Sneppen designed the study, performed the research and wrote the manuscript. Xiaochan Xu implemented models and all the simulations. All authors reviewed the manuscript.

## Data availability

The datasets generated during and/or analysed during the current study are available from the corresponding author on reasonable request.

